# The evolution of cooperation in the unidirectional linear division of labour of finite roles

**DOI:** 10.1101/2022.07.17.500384

**Authors:** Md Sams Afif Nirjhor, Mayuko Nakamaru

## Abstract

Evolution of cooperation is a puzzle in evolutionary biology and social sciences. Previous studies assumed that players are equal and have symmetric relationships. In our society, players are in different roles, have an asymmetric relationship, and cooperate together. We focused on the linear division of labour in a unidirectional chain that has finite roles, each of which is assigned to one group with cooperators and defectors. A cooperator in an upstream group produces and modifies a product, paying a cost of cooperation, and hands it to a player in a downstream group who obtains the benefit from the product. If players in all roles cooperate, a final product can be completed. However, if a player in a group chooses defection, the division of labour stops, the final product cannot be completed, and all players in all roles suffer damage. By using the replicator equations of the asymmetric game, we investigate which sanction system promotes the evolution of cooperation in the division of labour. We find that not the benefit of the product but the cost of cooperation matters to the evolutionary dynamics and that the probability of finding a defector determines which sanction system promotes the evolution of cooperation.

## 1 Introduction

Organisms including humans, plants, and bacteria cooperate (Dudley, 2015; Sachs and Hollowell, 2012; Boyd and Richerson 2009). Especially, human society is built on large-scale cooperation (Turchin, 2016; Harari, 2011). In the modern world, humans can often work with any other human from around the globe, disregarding country, ethnicity, language, and genetics. However, the evolution of cooperation in human society has remained a puzzle to unveil fully.

The prisoner’s dilemma (hereafter, PD game), which was first named by mathematician Albert Tucker, can describe why individuals fail to cooperate; if both are cooperators, their payoff is *R*; if both are defectors, their payoff is *P*. The payoff of a defector playing the PD game with a cooperator (*T*) is higher than the cooperator (*S*), and the order of the payoffs is *T > R > P > S*. As a result, players tend to choose defection rather than cooperation regardless of the opponent’s choice. Then, this game has its solution in mutual defection, although mutual cooperation would have given a better payoff for both. In other words, individual rationality is creating collective irrationality. This is called social dilemma (Kollock, 1998). However, cooperation exists in our society. What mechanism promotes the evolution of cooperation?

There are five possible mechanisms for creating cooperation in social dilemmas naturally without sanction, according to Nowak (2006) and Rand and Nowak (2013). They are kin selection (Hamilton, 1964), direct reciprocity (Axelrod and Hamilton, 1981; Veelen et al., 2012), indirect reciprocity (Sugden, 1986; Nowak and Sigmund 1998, 2005; Nakamaru and Kawata, 2004; Ohtsuki and Iwasa 2004), network reciprocity (Nowak and May, 1992; Nakamaru et al., 1997, 1998; Ohtsuki et. al., 2006; Pacheco et al., 2008; Santos et al., 2008) and group selection (Sober and Wilson, 1999; Traulsen and Nowak, 2006). Apart from these, using sanction or punishment to create cooperation have been studied (Axelrod, 1986; Sigmund et al., 2001; Boyd et al., 2003; Nakamaru and Iwasa, 2005, 2006; Rand et al., 2010; Sigmund et al., 2010; Sigmund et al. 2007; Shimao and Nakamaru, 2013; Chen et al., 2014, 2015; Sasaki et al., 2015).

Many previous theoretical studies about the evolution of cooperation considered that two or more players stand on the same footing. Rather, in reality, players take different positions, which the asymmetric game can describe. For an instance, parental investment to care their offspring was analyzed by the asymmetric game from the viewpoint of evolutionary game theory (Maynard Smith, 1977). It is because the fitness of the male is different from the female in parental care because the female reproduces egg but the male does not. This asymmetry causes sexual role, depending on the survivorship, the number of eggs and the chance to mate with other females. Another example of the asymmetric relationship among players is cooperation among different social roles where hierarchical social relationships exist (Henrich and Boyd, 2008; Powers and Lehmann, 2014; Roithmayr et al., 2015).

The division of labour is one of other examples of the asymmetric relationship between players (Kuhn and Stiner, 2006; Henrich and Boyd, 2008; Nakahashi and Feldman, 2014). In the division of labour, one player does not complete the whole role alone, but the whole role is divided among many smaller roles and each player is allocated to one role or more, each player only completes his/her role, and all players’ collaboration leads to the completion of the whole role. The division of labour can be observed not only in our human society but also in animals, plants, and bacteria (Iwasa and Yamaguchi, 2020; Wang et al., 2011; Dal et al., 2018).

In our human society, the historical and ethnographic evidence of division of labour’s presence in many pre-industrial societies and being associated with their development was shown in the previous studies (e.g. Nolan and Lenski, 2011; Durkheim, 1893). In our present world, there are many examples of division of labour, one of which is the division of labour by gender (Nakahashi and Feldman, 2014). The other example is supply chains observed in any industry (e.g. Fengru and Guitang, 2019). It also emerged in societies without centralized institutions such as governments (Chauvin and Ozak, 2017). Here comes the need for cooperation in the division of labour. However, the division of labour is a rather unexplored field in evolutionary game theory, especially when there are three or more roles in a system.

Nakamaru et al.(2018) explored the linear division of labour, in which roles are in line linearly, using the evolutionary game theory. They take the industrial dumping system in Japan as an example, in which cooperation and defection mean legal treatment and illegal dumping. Nakamaru et al.(2018) considered three roles, made concrete assumptions to fit the real industrial dumping system, and investigated if either of two existing sanction systems, namely the producer responsibility system and actor responsibility system, can promote the evolution of cooperation by means of the replicator equations for asymmetric games. In the former system if defection happens in the linear chain, whoever defects, the player in the first group gets punished by the supervision. In the later system, if defection happens, the defector is detected and gets punished by the supervision. It was shown that the sanction systems, especially the producer responsibility system when it is almost impossible to monitor and detect defectors, can promote cooperation more than the actor responsibility system. Hereafter, the former sanction system is called the first-role sanction system; the latter, is the defector sanction system.

In this study, we generalize the three role model into any countable number of roles model assuming a very simple model, and see whether the sanction systems can promote the evolution of cooperation or not.

## 2 Three Models

### 2.1 Baseline system

We present the baseline system (see figure 1), where there are *n* roles (*n ∈* **N** and *n ≥* 2). For *n* = 1, there is no linear division of labour. There are *n* groups in the whole population. Each group is allocated to one role, and the group size is infinite. Each group consists of cooperators and defectors. The frequency of cooperators in group *i* is *i*_*c*_ and the frequency of defectors is *i*_*d*_. Here, *i*_*c*_ + *i*_*d*_ = 1. It is assumed that one player chosen randomly from group *i* interacts with a player chosen randomly from group *i* + 1 (1 *≤ i < n*).

**Figure 1:**
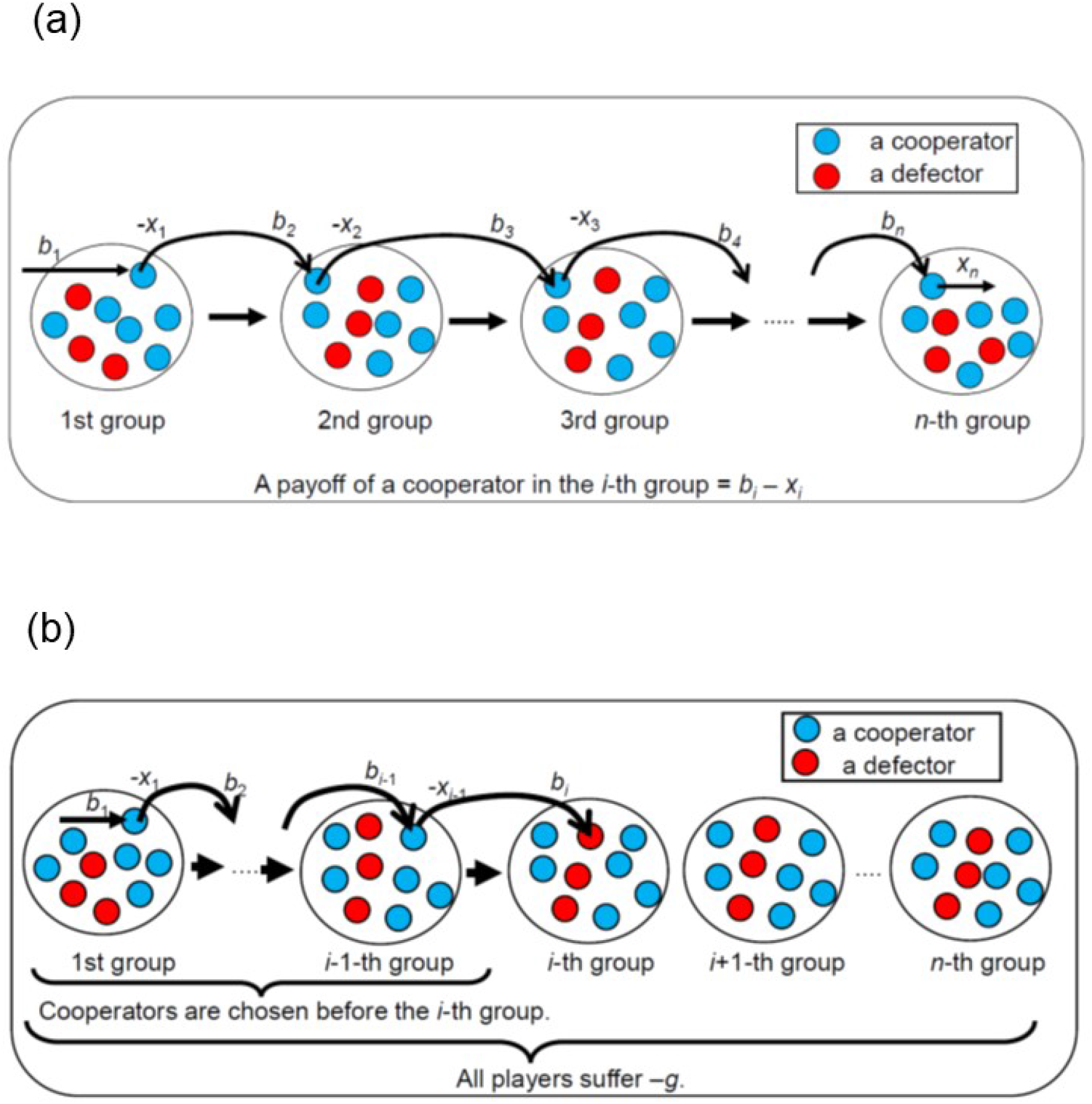
(a) Linear graph showing n division of labour when cooperators are chosen in all groups. The payoff of a cooperator in the *i*th group is *x*_*i−*1_–*x*_*i*_. (b) Linear graph showing n division of labour when cooperators are chosen before the *i*th group and then a defector is chosen in the *i*th group. Once a defector is chosen, the whole system is broken. As a result, all players suffer –*g*.

We consider that the final product or service is produced through the division of labour; if players in all roles cooperate together to produce the product or service, the final product or service can be completed (figure 1a). A cooperator in an upstream group produces and modifies the product or service, paying a cost of cooperation, and gives it to a player in a downstream group. Let *x*_*i*_ be defined as a cost of cooperation by a cooperator in group *i*. The value of the product or service is regarded as the benefit to the player in group *i* + 1, *b*_*i*+1_. In this case, the net benefit of cooperators in group *i* is *b*_*i*_ *− x*_*i*_ which should be non-negative (Tables 1a, b and c).

**Table 1a:** The payoff matrix for group 1 in the baseline system

| Cases | All being cooperator except the 1st | A defector after the 1st |
| --- | --- | --- |
| 1st player cooperator | $b_1 - x_1$ | $b_1 - x_1 - g$ |
| 1st player defector | $b_1 - g$ | $b_1 - g$ |

**Table 1b:** The payoff matrix in the baseline system for 1 *< i < n*

| Cases | All being cooperator except the $i$th | A defector before $i$th | A defector after $i$th |
| --- | --- | --- | --- |
| $i$th player cooperator | $b_i - x_i$ | $-g$ | $b_i - x_i - g$ |
| $i$th player defector | $b_i - g$ | $-g$ | $b_i - g$ |

**Table 1c:** The payoff matrix for group *n* in the baseline system

| Cases | All being cooperator except the $n$th | A defector before the $n$th |
| --- | --- | --- |
| $n$th player cooperator | $b_n - x_n$ | $-g$ |
| $n$th player defector | $b_n - g$ | $-g$ |

For the player in the 1st group, the benefit comes from the source. The net benefit of the cooperator in group 1 is *b*_1_ *− x*_1_ (Table 1a). In the *n*th group, a cooperator pays a cost, *x*_*n*_, to produce the final product. and the payoff of the cooperator is *b*_*n*_ *− x*_*n*_, where the player sells the final product and gets benefit *b*_*n*_ (Table 1c).

If a defector is chosen randomly from group *i*, the defector receives the benefit *b*_*i*_ and s/he does not cooperate with a player in group *i* + 1 (figure 1b). As a result, the division of labour fails and the whole system is broken down. As a result, all players in all the groups get the same negative effect *g* in their payoff. The payoff of the defector is *b*_*i*_ *− g* (Table 1a, 1b and 1c). After the defector is chosen from group *i*, the players in the latter groups will not choose either cooperation or defection, and there can only be one acting defector player in the chain of the linear division of labour here. Therefore, the net payoff of players after defection occurs is *−g* (Tables 1b and c).

Table 1 indicates that, when *g < x*_*i*_, the game can be the PD game. Otherwise, mutual cooperation is always the best. Therefore, we assume that *g < x*_*i*_, which means that the baseline system has the dilemma situation.

For calculating player *i*’s payoff matrix we need to consider three cases. First, when there are all cooperators in the chain of division except group *i*, the probability is given by *c*_*i*_, where, 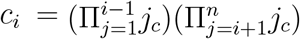 Second, when there is a defector before group *i*, it’s probability is given by *d*_*ib*_,.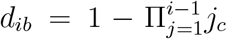 Finally, when there is a defector after *i*, its probability is given by *d*_*ia*_, where 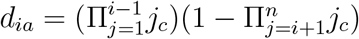. Here, *c*_*i*_ + *d*_*ib*_ + *d*_*ia*_ = 1.

After each player randomly chosen from each group interacts with a player randomly chosen from the next downstream group, the expected payoff of each player in each group can be calculated. Then, within each group, players decided to imitate a strategy of others, proportional to the expected payoff relative to the total payoff in the group. Here the random change of the strategy or mutation does not occur. Therefore, this interaction can be described by the replicator dynamics of asymmetric game without mutation (Hofbauer and Sigmund, 1998).

The replicator equation of a cooperator in group 1 in the baseline system is as following,

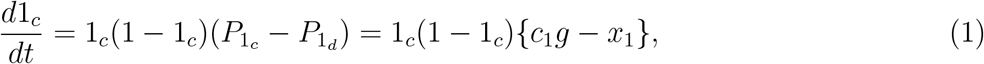

where the average payoff of cooperator in group 1, 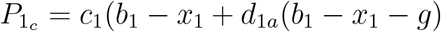 and the average payoff of the defector in group 1, 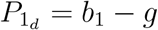.

The replicator equation for cooperators in group 1 *< i < n* is as following,

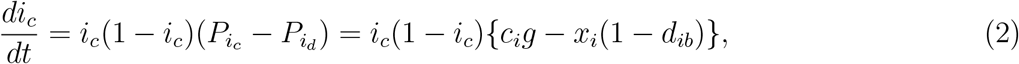

where the average payoff of cooperator in group 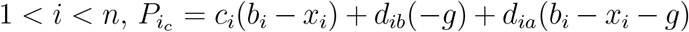 and the average payoff of the defector in group 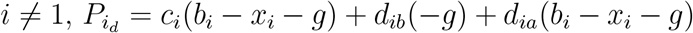.

The replicator equation for cooperators in group *n* is as following,

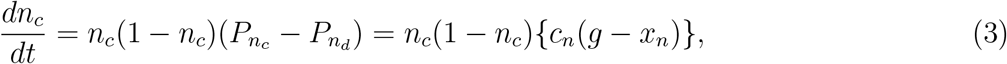

where the average payoff of cooperator in group *n*, 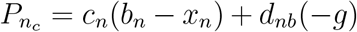 and the average payoff of the defector in group *n*, 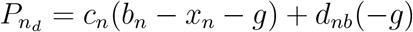.

### 2.2 The defector sanction system

Next we focus on two sanction systems, namely the defector sanction system, and the first role sanction system. In the defector sanction system, the defector in the chain of the linear division of labour gets punished with a fine *f*, where (*f >* 0) and the finding probability of the defector is *ρ*. In some types of linear division of labour, where monitoring a defector is too hard, where *ρ* is very low compared to other parameters.

For the defector sanction system, the payoffs are given in a similar way as the baseline except the punishment; adding the punishment of *ρf* to the defector’s payoff. In the defector sanction system, the payoff matrix for player 1 is presented in Table 2a, the payoff matrix for a player in group 1 *< i < n* is in Table 2b and the payoff matrix for a player in group *n* is in Table 2c.

**Table 2a:**
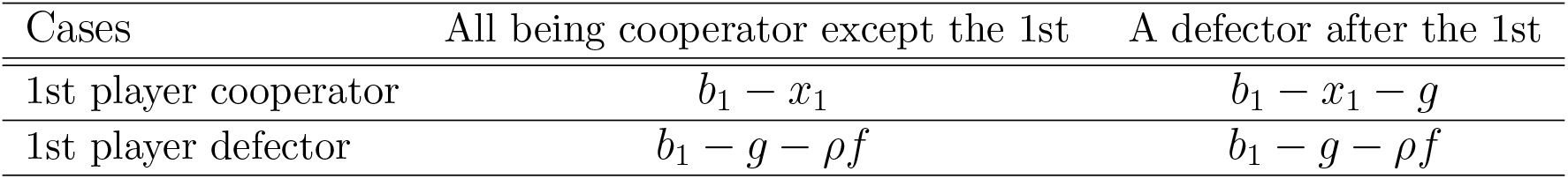
The payoff matrix in the defector sanction system for group 1

| Cases | All being cooperator except the 1st | A defector after the 1st |
| --- | --- | --- |
| 1st player cooperator | $b_1 - x_1$ | $b_1 - x_1 - g$ |
| 1st player defector | $b_1 - g - \rho f$ | $b_1 - g - \rho f$ |

**Table 2b:** The payoff matrix in the defector sanction system for 1 *< i < n*

| Cases | All being cooperator except the $i$th | A defector before $i$th | A defector after $i$th |
| --- | --- | --- | --- |
| $i$th player cooperator | $b_i - x_i$ | $-g$ | $b_i - x_i - g$ |
| $i$th player defector | $b_i - g - \rho f$ | $-g$ | $b_i - g - \rho f$ |

**Table 2c:** The payoff matrix in the defector sanction system for group *n*

| Cases | All being cooperator except the $n$th | A defector before the $n$th |
| --- | --- | --- |
| $n$th player cooperator | $b_n - x_n$ | $-g$ |
| $n$th player defector | $b_n - g - \rho f$ | $-g$ |

The replicator equation for the cooperators in group 1 is as following,

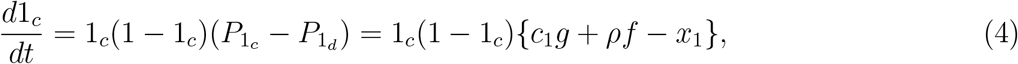

where the average payoff of cooperator in group 1, 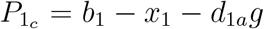 and the average payoff of the defector in group 1, 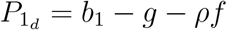.

In the defector sanction system, replicator equation for the cooperators in group 1 *< i < n* is as following,

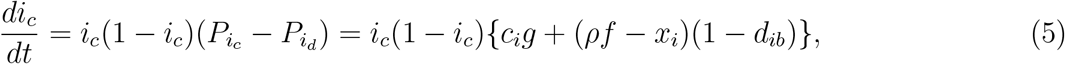

where the average payoff of cooperator in group 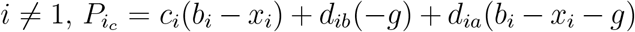 and the average payoff of the defector in group 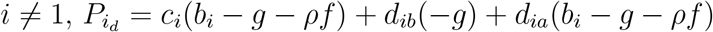.

In the defector sanction system, the replicator equation for the cooperators in group *n* is as following,

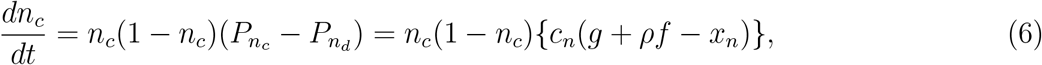

where the average payoff of cooperator in group *n*, 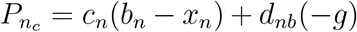 and the average payoff of the defector in group *n*, 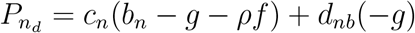.

### 2.3 The first role sanction system

In the first role sanction system, the fine is the same as the defector sanction system, *f*, and a defection in the chain of the linear division of labour is always found with probability 1, because the final product or service does not appear if defection occurs, and players can know that defection occurs without monitoring a defector. Therefore, the finding probability is one. The player in the first role always gets punished for the defection, no matter which role defected. For example, this sanction system is executed to prevent illegal dumping in Japan (Nakamaru et al., 2018).

For the first role sanction system for the generalized *i*th player the payoff matrix is the same as the baseline except that for group 1. If anyone defects, the first role gets punished, and punishment *f* appears in the first group’s payoff matrix (Table 3a). The payoff matrix for a player in the group 1 in the first role sanction system is in Table 3a, for a player in the group 1 *< i < n*) in the first role sanction system is in Table 3b and for a player in group *n* is in Table 3c.

**Table 3a:**
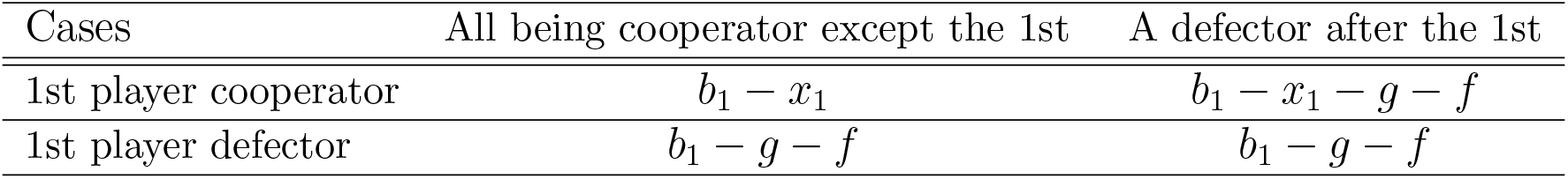
The payoff matrix in First role sanction system for group 1

**Table 3b:** The payoff matrix in first role sanction system for *i /*= 1

| Cases | All being cooperator except the $i$th | A defector before $i$th | A defector after $i$th |
| --- | --- | --- | --- |
| $i$th player cooperator | $b_i - x_i$ | $-g$ | $b_i - x_i - g$ |
| $i$th player defector | $b_i - g$ | $-g$ | $b_i - g$ |

**Table 3c:**
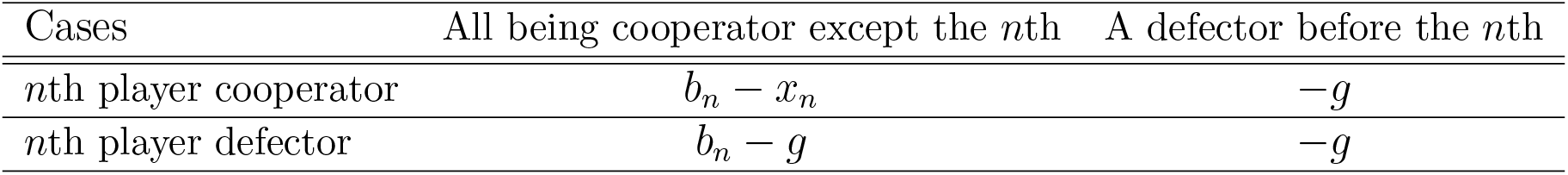
The payoff matrix for group *n* in the baseline system

The replicator equation for a cooperator in the group 1 is as following;

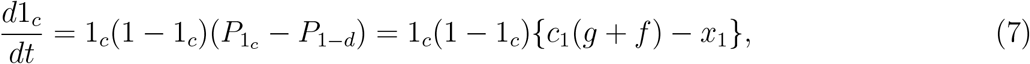

where the average payoff of cooperator in group 1, 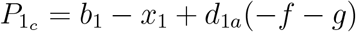 and the average payoff of the defector in group 1, 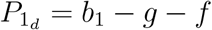.

The replicator equation for a cooperator in the group 1 *< i < n* and group *n* are the same as the baseline model. The replicator equation for a cooperator in the group 1 *< i < n* is

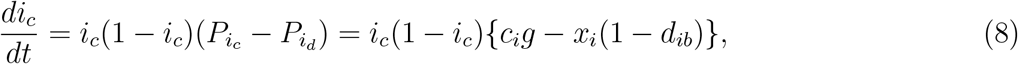

where the average payoff of cooperator in group 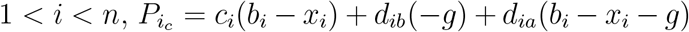 and the average payoff of the defector in group 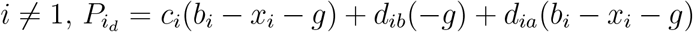.

The replicator equation for cooperators in group *n* is as following,

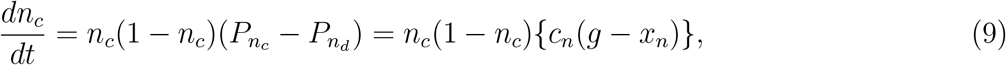

where the average payoff of cooperator in group *n*, 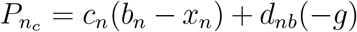 and the average payoff of the defector in group *n*, 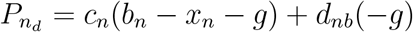.

Table 4 shows the parameter list in our model.

**Table 4:** Parameters

|  |  |
| --- | --- |
| $n$ | The number of groups, ($n \in \mathbf{N}$ and $n \geq 2$) |
| $i$ | Index of groups, $1 \leq i \leq n$ |
| $b_i$ | The benefit given by the $i$th group player to the $i + 1$th group player |
| $x_i$ | The cost of cooperation for the $i$th group player |
| $i_c$ | The frequency of cooperators in the $i$th group |
| $i_d$ | The frequency of defectors in the $i$th group |
| $g$ | A damage for all players once one defector appears in the line |
| $f$ | The amount of punishment |
| $\rho$ | The probability of catching a defector |
| $c_i$ | The probability of having all cooperators in the line except the $i$th group |
| $d_{ib}$ | The probability of having a defector in the line before the $i$th group |
| $d_{ia}$ | The probability of having a defector in the line after the $i$th group |

## 3 Results

### 3.1 The summary of results in all three systems

We find three sorts of equilibrium in all three systems. One is all cooperating which we represent as [1_*c*_, 2_*c*_, …, *n*_*c*_] = [1, 1, …, 1], then all defecting which we represent as [1_*c*_, 2_*c*_, …, *n*_*c*_] = [0, 0, …, 0], which is also represented by [0, *∗*, …, *∗*] where ”*∗*” is any value between 0 and 1. Because the game stops there and the players in the later roles gets the same payoff regardless of the behaviour once the player in group 1 defects. As a result, *i*_*c*_ neutrally converges to any value between 0 and 1 (*i >* 1). The third one is cooperating-defecting mixed equilibrium which is represented as [1_*c*_, 2_*c*_, …, (*j −* 1)_*c*_, *j*_*c*_, …, *n*_*c*_] = [1, 1, …, 1, 0, …, 0], where *j* is between 1 and *n −* 1. It can be rewritten as [1, 1, …, 1, 0, *∗*…, *∗*]. We analyse local stability of these three equilibria with the Jacobian matrix applying Routh-Hurwitz criteria (see Appendix). Table 5 presents the summary of the analyses.

**Table 5:** Local stability conditions for the general model

| Equilibrium | Baseline | Defector sanction | First role sanction |
| --- | --- | --- | --- |
| $[0, \dots, 0]$ | Always stable | $\rho f < x_1$ | Always stable |
| $[1, \dots, 1, 0, *, \dots, *]$ | Always unstable | $\max\{x_i\}_{i=1}^{j-1} < \rho f < x_j(*)$ | Always unstable |
| $[1, \dots, 1]$ | $g > \max\{x_i\}_{i=1}^n$ | $\rho f + g > \max\{x_i\}_{i=1}^n$ | $f + g > x_1$ & $g > \max\{x_i\}_{i=2}^n$ |

(\*): $j$ is the first defector

Table 5 shows that the all defection equilibrium is one stable equilibrium in the baseline. It is also stable in the first role sanction system. However, it is stable in the defector sanction system, if *ρf > x*_1_.

The mixed equilibrium is unstable in both the baseline and the first role sanction system. It is stable in the defector sanction system when *ρf < x*_*j*_ and 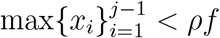, where player *j* defects and everyone cooperates before player *j*.

The all cooperation equilibrium is locally stable if 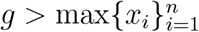 in the baseline system. However, we consider the PD situation in the baseline system, and then we can assume that *g < x*_*i*_. This indicate that all cooperation is unstable in the baseline. It is stable in the defector sanction system when 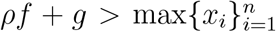, which is held even though *g < x*_*i*_. Therefore, if *ρf* is large enough, the defector sanction system can promote the evolution of cooperation.

The all cooperation equilibrium is stable in the first role sanction system when *f* + *g > x*_1_ and 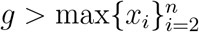. This indicates that the first role sanction system promotes the evolution of cooperation more than the baseline system. If we consider the assumption, *g < x*_*i*_, which satisfies the condition of the PD game, the all cooperation equilibrium is considered unstable in the first role sanction system.

Appendix and Table 5 suggest that the benefit given by the *i*th group player to the *i* + 1 th group player, *b*_*i*_, does not influence the local stability of each equilibrium point.

To understand the dynamics well, we will discuss three special cases about the cost of cooperation in the following sections.

### 3.2 Three special cases

#### 3.2.1 The cost of cooperation is lower in higher *i*

Here, we consider the special case where the cost in a downstream group decreases in the linear division of labour, *x*_1_ *> x*_2_ > …‥ > *x*_*n*_.

After exploring the local stability conditions for each of the equilibrium in each of three systems, we can summarize the results as Table 6, which represents that the all defection equilibrium is a stable equilibrium in the baseline, and is also stable in the first role sanction system. In the defector sanction system, it is locally stable when *ρf < x*_1_.

**Table 6:**
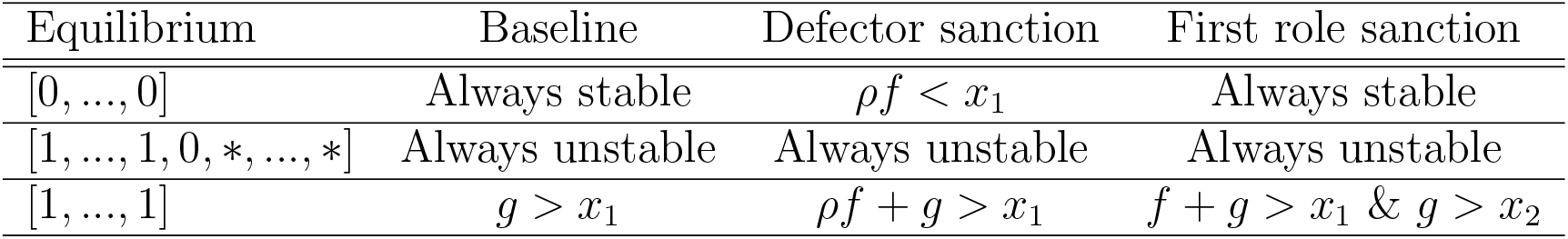
Local stability conditions when the cooperation cost decreases in the downstream

| Equilibrium | Baseline | Defector sanction | First role sanction |
| --- | --- | --- | --- |
| $[0, \dots, 0]$ | Always stable | $\rho f < x_1$ | Always stable |
| $[1, \dots, 1, 0, *, \dots, *]$ | Always unstable | Always unstable | Always unstable |
| $[1, \dots, 1]$ | $g > x_1$ | $\rho f + g > x_1$ | $f + g > x_1$ & $g > x_2$ |

The mixed equilibrium is unstable in all the system. This indicates that once cooperation in group 1 starts, the all cooperation equilibrium will be stable. It is intuitive because all players face the same cost *−g* once a player chooses defection and the cost of cooperation *x*_*i*_ is lower in the later groups. Then, the players in the later groups will follow the cooperator. It is meaningful to punish a defector in the earliest group to promote cooperation. Therefore, the first role sanction system works.

Our analysis suggests that the all cooperation equilibrium is stable in the baseline system only when *g > x*_1_ because *x*_1_ *> x*_2_ *>* ….. *> x*_*n*_. As we assume that *g < x*_1_ which meets the PD game, the equilibrium is not stable (figure 2(a)). It is stable when *ρf* + *g > x*_1_ for the defector sanction systems (figure 2(b)), and is stable when *f* + *g > x*_1_ and *g > x*_2_ for first role sanction system (figure 2(c)). Figure 4(d) shows that the same punishment *f* as the first role sanction system cannot create evolution of cooperation in the defector sanction system, because of the low finding probability of the defector.

**Figure 2:**
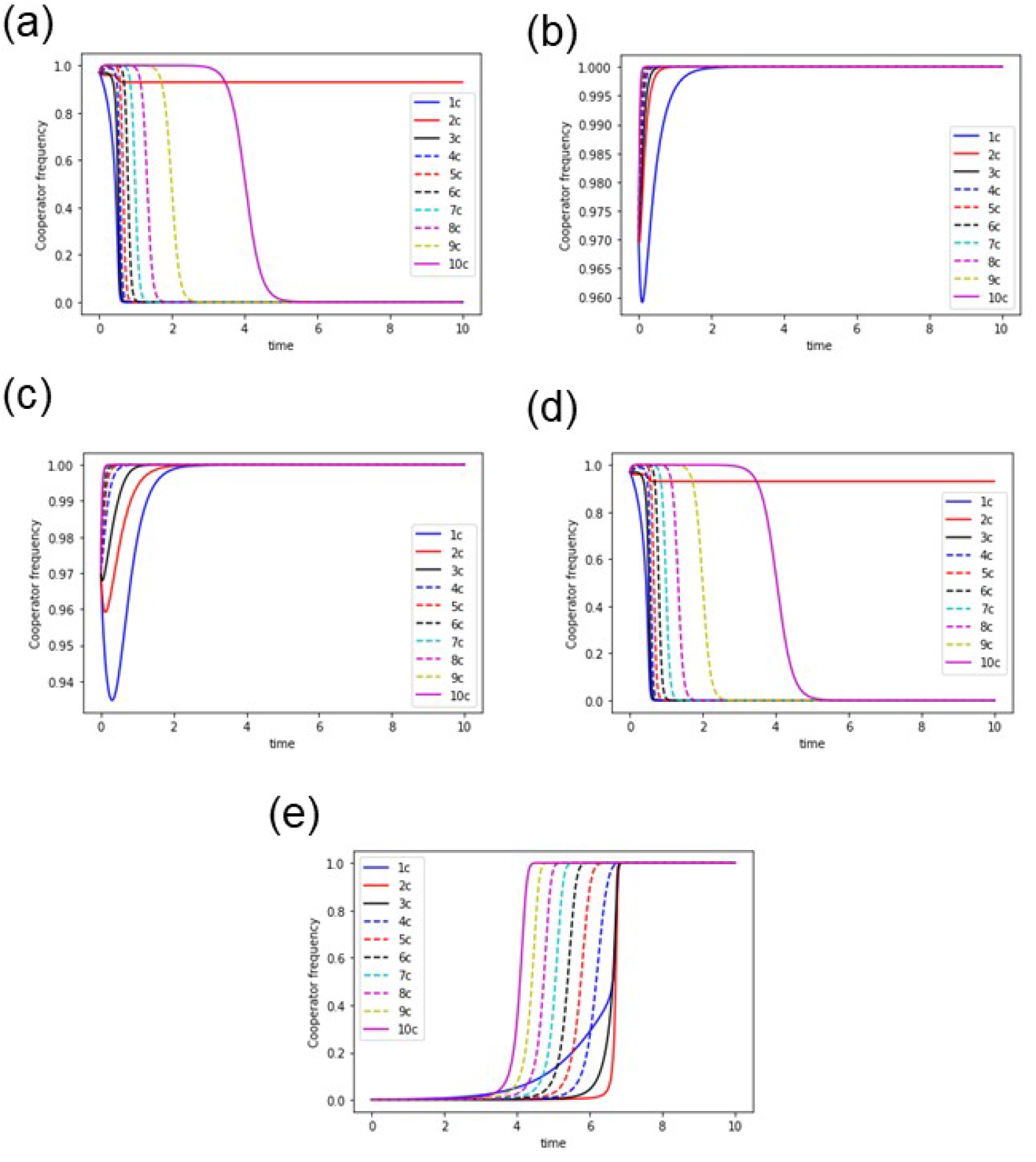
Dynamics of the system with *n* = 10 when cooperation cost decreases downstream. (a) The baseline, (b) The defector sanction system with punishment *f* = 5000, making *ρf* = 5, (c) The first role sanction system with punishment *f* = 5. This shows that when *i*_*c*_(0) is high, defector sanction system needs higher amount of punishment *f* than the first role sanction system because of its dependence on the finding probability of the defector, *ρ*. (d) Defector sanction system with punishment *f* = 5, making *ρf* = 0.005. (e) The defector sanction system has *ρf* = 51. Even when the initial frequency of cooperators are rare, the evolution of cooperation happens if *ρf > x*_1_. The parameters are: *i*_*c*_(0) = 0.97 for (a), (b), (c) and (d); *i*_*c*_(0) = 0.001 for (e); *g* = 48, *ρ* = 0.001, *x*_1_ = 50 and *x*_*i−*1_ *− x*_*i*_ = 5 (for all *i*s) for (a), (b), (c), (d), and (e).

Figures 2(e) and 3(a) show the defector sanction system promotes cooperation even though cooperators are rare in the beginning in *ρf > x*_1_. In the region where *ρf* + *g > x*_1_ and *ρf < x*_1_, the system is bistable, where the punishment needs to be higher to create all cooperation with lower *i*_*c*_(0), and even low punishment can create all cooperation with higher *i*_*c*_(0). Figure 3(a) also shows that the dynamics is independent of the initial condition and goes to all defection in the region of *ρf* + *g < x*_1_. Therefore, if the probability of finding and catching a defector is too low and *ρf* is very low, the defector sanction system never promotes cooperation. If the punishment, *ρf*, is large enough to be effective, the defector sanction system promotes cooperation even though the initial frequency of cooperators is low.

**Figure 3:**
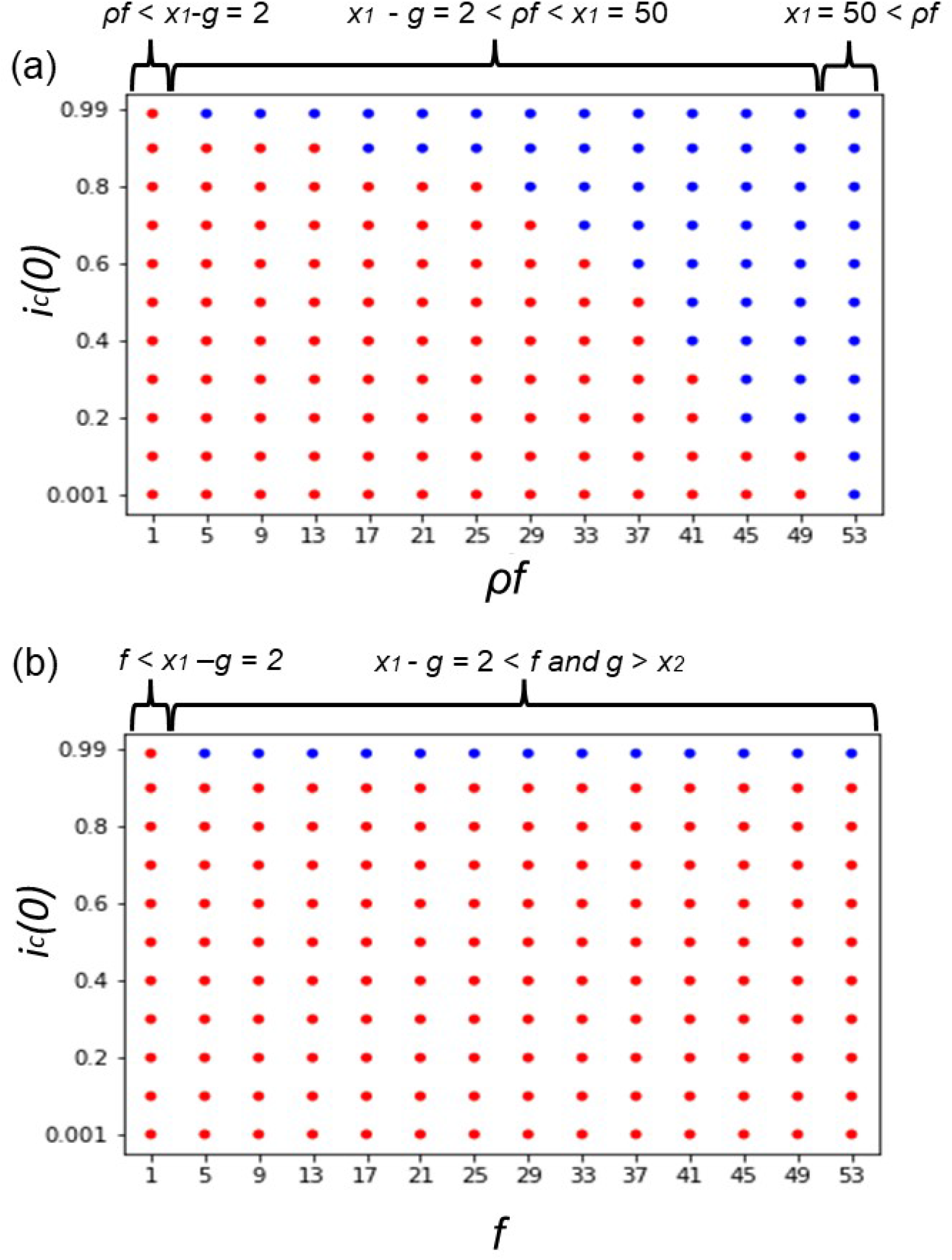
Initial frequency dependency in (a) the defector sanction system and (b)the first role sanction system with *n* = 10 when the cooperation cost decreases downstream. The horizontal axis is for *ρf* in (a) and for *f* in (b). The vertical is for the initial frequency of cooperators in group *i, i*_*c*_(0), when *i* is an integer between 1 and 10. Blue dots shows when the dynamics evolves into all cooperation and red dots shows when the dynamics evolves into all defection. The parameters are: *g* = 48, *x*_1_ = 50, *ρ* = 0.001 and *x*_*i−*1_ *− x*_*i*_ = 5 for all *i*s.

Figure 3(b) shows clearly that the first role sanction system creates cooperation only when the initial frequency of cooperators in all the groups are very high. When the *i*_*c*_(0) comes near 0.95, cooperation only evolve with very high punishment *f* (figure 2(c)). When *i*_*c*_(0) is low, the system goes to all defection.

#### 3.2.2 The cost of cooperation is the same for all the groups

We assume *x*_*i*_ is the same for all the 1 *≤ i ≤ n*; *x*_1_ = … = *x*_*n*_ = *x*. After exploring the local stability conditions for each of the equilibrium in each of three systems (see appendix), we can summarize the results as Table 7, which are basically the same as Table 6.

**Table 7:**
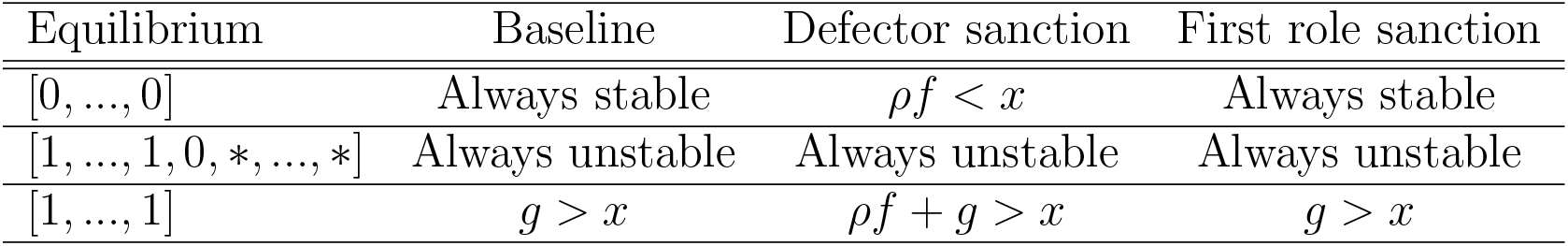
Local stability conditions when the cooperation cost is the same for all the groups

#### 3.2.3 The cost of cooperation is higher in higher *i*

Now we consider the model when the cost of cooperation rises in the downstream of the linear chain. Here we assume that, *x*_*i*_ *> x*_*i−*1_ for all *i*s. The local stability conditions for each of the three equilibria are shown in table 8.

**Table 8:** Local stability conditions when the cooperation cost increases in downstream

| Equilibrium | Baseline | Defector sanction | First role sanction |
| --- | --- | --- | --- |
| $[0, \dots, 0]$ | Always stable | $\rho f < x_1$ | Always stable |
| $[1, \dots, 1, 0, *, \dots, *]$ | Always unstable | $x_{j-1} < \rho f < x_j(*)$ | Always Unstable |
| $[1, \dots, 1]$ | $g > x_n$ | $\rho f + g > x_n$ | $g > x_n$ |

(\*): $j$ is the first defector

The main difference between the condition where the cost of cooperation increases in downstream groups and the condition where the cost of cooperation decreases in downstream groups is that the mixed equilibrium can be stable when the cost of cooperation increases in downstream groups (Table 8).

Figure 4(a) shows that the dynamics is completely independent of the initial frequency of cooperators in the defector sanction system; in the region of *ρf < x*_1_ the dynamics goes to all defection, even with very high initial frequency of cooperators in all groups (figure 5(a)). When *x*_*j−*1_ *< ρf < x*_*j*_, all players cooperates till the *j −* 1th group before a player in the *j*th group chooses defects; we get the mixed equilibrium shown in figure 5(b) and 5(c). When *ρf* + *g > x*_*n*_, all players in all groups go to cooperation even though their initial frequencies of cooperators are small (figure 5(d)).

**Figure 4:**
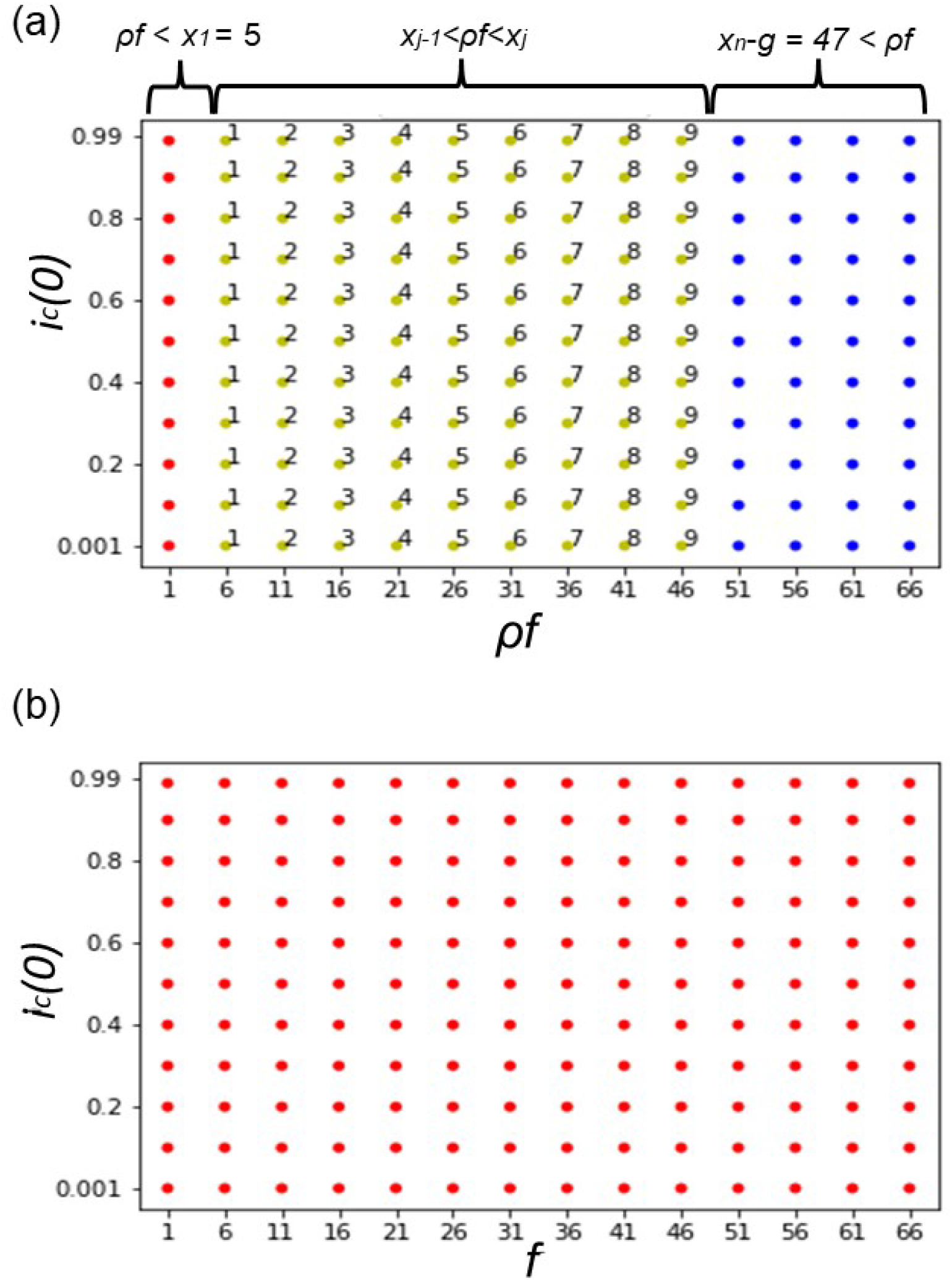
Initial frequency dependency in (a) the defector sanction system and (b) the first role sanction system with *n* = 10 when the cooperation cost increases downstream. The horizontal axis is for *ρf* in (a) and for *f* in (b). The vertical axis is for *i*_*c*_(0) when *i ∈ {*1, 2, 3, 4, 5, 6, 7, 8, 9, 10*}*. The parameters are: *g* = 3, *x*_1_ = 5. *ρ* = 0.001 and *x*_*i*_ *− x*_*i−*1_ = 5 for all *i*s. Blue dots show when the dynamics evolves into all cooperation and red dots shows when the dynamics evolves into all defection. The yellow dots in (a) show when the dynamics evolves into a mixed equilibrium. The number outside each yellow dot shows which final group has the all cooperative population in that mixed equilibrium, or *j −* 1. As *x*_*j−*1_ *< ρf < x*_*j*_ is the condition for the mixed equilibrium to be stable where *j* is the first defector.

**Figure 5:**
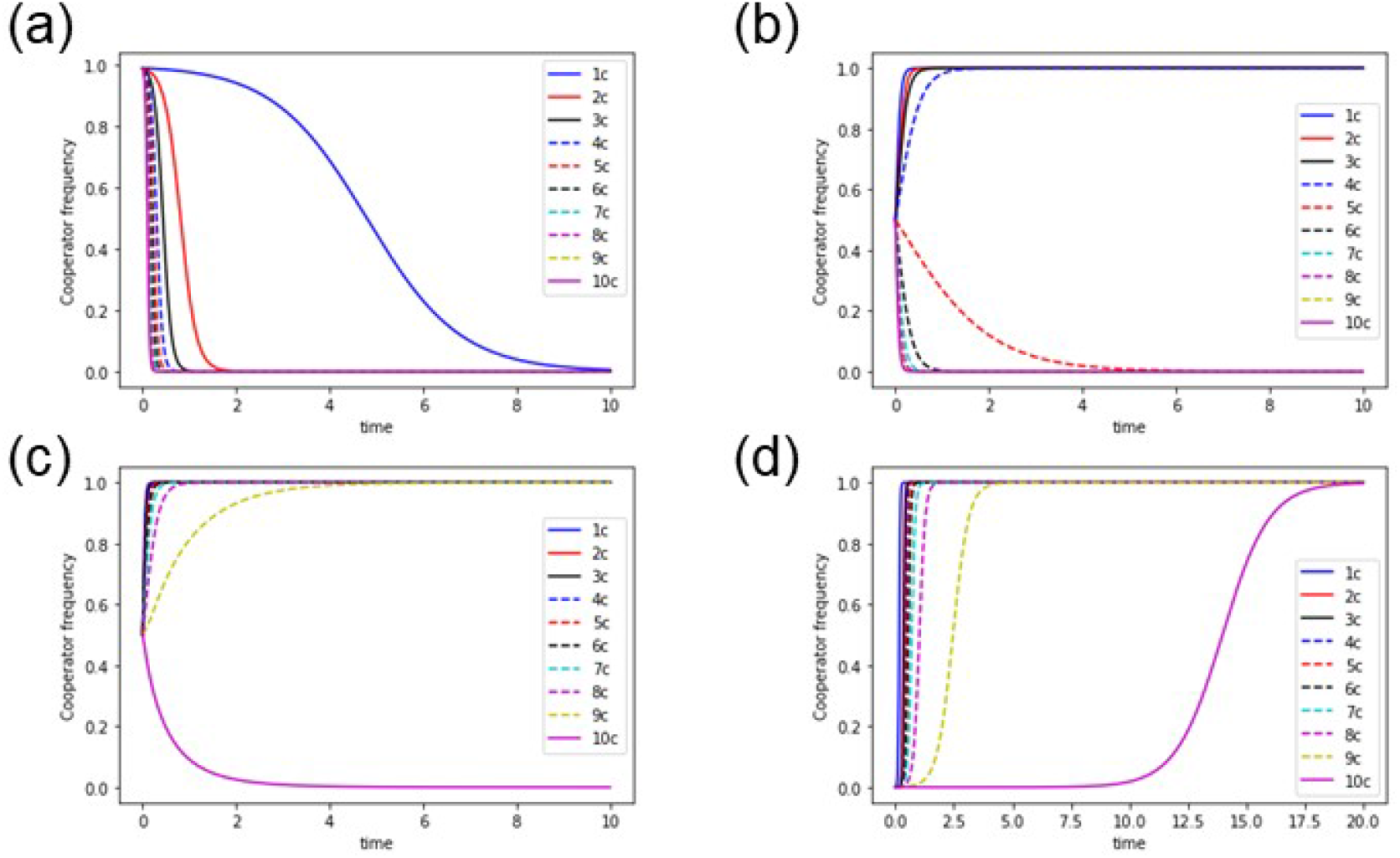
Time evolution in the defector sanction system with *n* = 10 when cooperation cost increases downstream. The horizontal axis is for time and the vertical one is for frequency of cooperators in each group. In (a) *ρf* = 4 and *i*_*c*_(0) = 0.99 shows even when the initial frequency of cooperators in all groups are very high, because of *ρf < x*_1_, all defection equilibrium is stable. In (b) *ρf* = 24, *i*_*c*_(0) = 0.5 shows the dynamics evolving into the mixed equilibrium where all players are cooperators until group 4 and in group 5 all are defectors as *x*_4_ *< ρf < x*_5_. In (c) *ρf* = 46, *i*_*c*_(0) = 0.5 shows the dynamics evolving into the mixed equilibrium where all are cooperators until group 9 and all are defectors in group 10, as *x*_9_ *< ρf < x*_10_. In (d) *ρf* = 48, *i*_*c*_(0) = 0.001 shows even when the initial frequency of cooperators in all groups are very low, because of *ρf > x*_*n*_ *− g*, the all cooperation equilibrium is stable. The parameters are: *g* = 3, *x*_1_ = 5, *ρ* = 0.001, and *x*_*i*_ *− x*_*i−*1_ = 5 for all *i*s.

Figure 4(b) shows that first role sanction system can never create all cooperation, even with very high punishments *f* and along all the initial cooperation frequency.

In sum, the defector sanction system works as sanction and promotes the evolution of cooperation when *x*_1_ *< x*_2_ *<*.. *< x*_*n*_ and *ρf* is large enough. While, the first role sanction system does not work and it is equivalent to the baseline system.

## 4 Discussion and Conclusion

We took a system of linear division of labour where there are *n* roles (*n ≥* 2). If a role gets subjected to defection by its defector, the labour stops there, and the players associated with the later roles do not get a chance to play the roles. Each player in each group gets subjected to the same loss once a player defects. We analyse three systems; the baseline system and the two sanction systems namely the defector sanction system and the first role sanction system, to see their effect on the evolution of cooperation. After applying the replicator equation of asymmetric game, we find three equilibria, 1) where all the players in all the groups are defectors, 2) where all the players in all the groups are cooperators and 3) where the players in the earlier groups are all cooperators and the players in the later groups are defectors which is called the mixed equilibrium.

Our findings are as follows: the benefit given by a cooperator in an upstream group to a player in a downstream group does not influence the evolutionary dynamics, but the cost of cooperation does. We compare two sanction system, the defector sanction system and the first role sanction system, with the baseline system. Then, we found that the defector sanction system promotes the evolution of cooperation unless the probability of finding a defector is very low. However, when it is too hard to monitor and detect a defector, the defector sanction system does not work as sanction anymore. The first role sanction system promotes cooperation when the cost of cooperation decreases in downstream groups. Otherwise, the first role sanction system is equivalent to the baseline system; it does not work as sanction. The other important point is that the mixed equilibrium can be locally stable when the cost of cooperation increases with higher *i*.

We have found a trend that (i) the frequency of cooperators in the group with a lower cost of cooperation reaches one faster when either the all cooperation equilibrium or the mixed equilibrium is locally stable (figures 2b, 2c, 2e, 5b, 5c, and 5d), and (ii) the frequency of defectors in the group with a higher cost of cooperation reaches one faster when either all defection equilibrium or the mixed equilibrium is stable (figure 2a, 2d, 5a, 5b and 5c). This shows the comparative effect of the cost of cooperation among different groups on the dynamics. We only did the numerical analysis when all of the groups are given the same initial condition, and this trend may be dependent on the initial condition. To investigate whether the initial condition changes this trend, more numerical analysis will be needed.

There are some evolutionary game theoretical studies that seem similar to our framework. The effect of network structures on creating cooperation in asymmetric social interactions has been studied (e.g. McAvoy and Hauert, 2015; Su et al., 2022). Their studies seem to include ours, however it is not true. In their studies, each player is in each vertex, a player imitates the strategy of the neighbour who is a partner of the game if the payoff is higher than that of the player. For example, Su et al. (2022) concentrated on how edge orientation, directionality, regularity and properties of graphs will change the evolution of cooperation. The cooperation cost and benefit for all players are the same. In our model, each group is in the line and has a large number of players. A player imitates the strategy of others in the group, and a player chosen randomly from an upstream group never play the game with another player in the same group but does with a player chosen randomly in the downstream group. Our study focused on the impact of sanctions in creating cooperation with a graphical structure of unidirectionality when there are a starting point and terminal point in the linear network. The players in different roles have the different cooperation cost and benefit.

Our study also might remind us of Boyd and Richerson (1989). In their study, the action of a player towards his downstream player depends on the action of either the upstream player towards him or the downstream player to his downstream player in a cycle network. Two neighbours in the unidirectional cycle network play the PD game in order and it goes on repeated. Among the two strategies, 1) Upstream tit for tat (UTFT): If the upstream player cooperates/defects, the player cooperates/defects with the downstream player, 2) Downstream tit for tat (DTFT): If the downstream player cooperates/defects with his downstream player, in the later cycle the player cooperates/defects with the downstream player, DTFT evolved under more circumstances than UTFT. Structurally this study might look similar to ours. However, there are some critical differences between the two. One difference is the research purpose; we investigated the evolution of cooperation in the linear division of labour on the basis of sanction systems; Boyd and Richerson (1989) investigated the evolution of indirect reciprocity. Our game structure is line; our study assumed each group is on each site and does not assume that players in two neighbouring groups interact repeatedly. While, the former study’s network structure is a repeated cycle and each player is on each site.

The studies about the evolution of cooperation assuming that each group is located at the two dimensional lattice, players move to their neighbouring groups (e.g. Wakano et al., 2009) also seem similar to ours. However, our study did not assume the movement of players to the neighbouring groups but the interaction between a player in a group and a player in a downstream group to describe the division of labour.

Therefore, we conclude that some previous studies which seem similar to our study are different from ours, and our study gives a new perspective that the division of labour can be studied from the viewpoint of “the evolution of cooperation.”

We consider the special case that *b*_*i*_ = *x*_*i−*1_, which means the benefit given by a cooperator in group *i −* 1 is same as the cost of cooperation paid by the cooperator. As *b*_*i*_–*x*_*i*_ = *x*_*i−*1_–*x*_*i*_ should be positive, *x*_*i*_ decreases as the *i* increases. Therefore, the result of the analyses corresponds to Table 6. The total sum of the net benefit of all cooperators in all roles is = (*x*_0_–*x*_1_) + (*x*_1_–*x*_2_) + + (*x*_*n−*1_–*x*_*n*_) = *x*_0_–*x*_*n*_. Therefore, the assumption that this situation can be interpreted as that each player in each role decides how much the upstream player keeps and distribute to the downstream players (figure 6). For example, a cooperator in group 1 keeps *x*_0_–*x*_1_ and gives *x*_1_ to a player in group 2. This situation has some implications for government planning and spending because the model can be assimilated with the flow of government spending (see figure 6). For government planning and spending, cooperation in the division of labour is required (Davis, 2001). Government of all forms performs a very significant role in running modern country states. The roles of it are as elaborated as they can get and a single individual cannot perform all roles. Therefore, all the related tasks are divided among many ministries and the ministries also divide the tasks among their workers. For a certain work to be done at the root level, the fund should go through multiple agents as well as be planned by multiple agents. Therefore, cooperation of all the roles is a key for success in that particular task. For example, if a governmental head wants to spend some money at the root level, he/she allocates the money to his/her subordinate, who also allocates the part of the money to his/her subordinate, and it goes to many levels of subordinates until reaching the goal. Every cooperator in the group *i* here gains the net benefit *x*_*i−*1_ *− x*_*i*_ from cooperation (see figure 6(a)). And if someone stops or does not cooperate because of corruption or other reasons, the money does not reach the goal and therefore the task fails. As a result, all players additionally get damage, *−g* (see figure 6(b)). In return, the defector, of course, gains the money which was given to him but to every agent as a part of the government comes a bad reputation for the defection which can be set as *−g*. By creating cooperation among all the roles, we make sure that the players in the first role engaging in government spending do not choose defection. The reason is as follows; our results suggest that all cooperation equilibrium or all defection equilibrium can be locally stable but the mixed equilibrium is not stable. This means if a player in the first group is a cooperator, players in all other groups can be cooperators.

**Figure 6:**
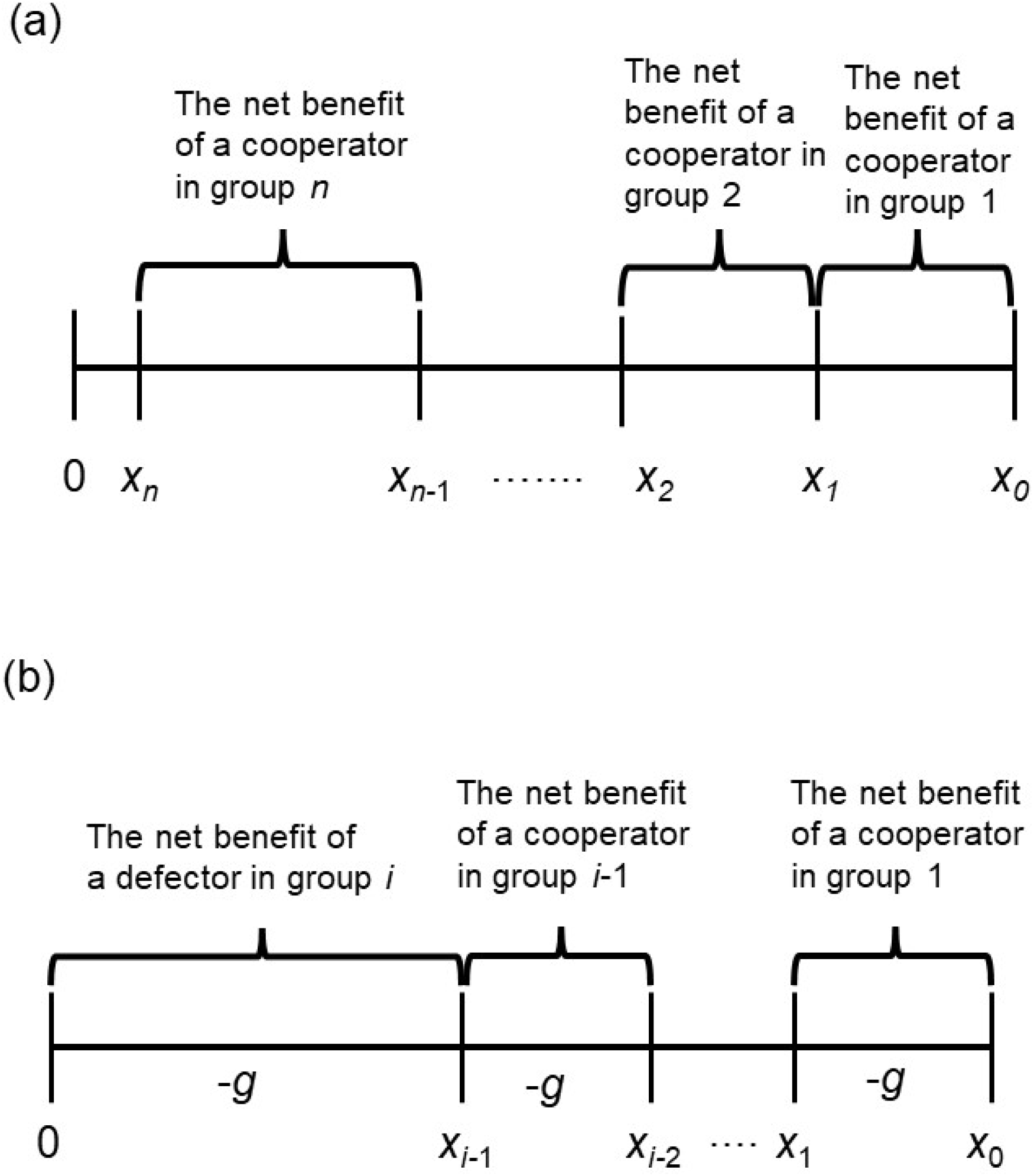
Allocation of the benefit among the players. Then net benefit of a player in group *i* is equal to the amount that the player keeps. (a) shows when all players in all groups are cooperators, (b) shows when there is a defector is chosen in the group *i*.

We did not comment particularly on the impact of the net benefit *b*_*i*_ *− x*_*i*_ or the distribution of the benefit for the cooperators in the system because the benefit does not influence the dynamics (see Tables 5-8). While the benefit from a cooperator is also cancelled out in the previous study about the division of labour in the industrial dumping system (Nakamaru et al., 2018). In Nakamaru et al. (2018) which made the concrete model fitting the industrial dumping system, the cost of cooperation decreases in the downstream roles. Nakamaru et al. (2018) concluded that the first role sanction system promotes cooperation more than the defector sanction system especially when the probability of finding a defector is very low, which is also supported by our result when the cost of cooperation decreases in downstream groups. The difference between these two studies is that there are locally stable mixed equilibria such as (1_*c*_, 2_*c*_, 3_*c*_) = (1, 1, 0) in the baseline and two sanction systems in Nakamaru et al. (2018), but in the defector sanction system when the cost of cooperation increases in downstream groups in this study. In our future research, we will investigate what causes the difference.

In this paper, we consider a very simple case in the linear division of labour. Furthermore, we will develop our model and make the model when there are general and complex networks in the division of labour.

## Acknowledgement

NMSA was supported by JST, the establishment of university fellowships towards the creation of science technology innovation, Grant Number JPMJFS2112. MN was supported by JSPS KAKENHI Grant Number JP21K01626.

## Appendix

We analysed the local stability of the equilibria in each of the three systems for the general model.

For the baseline system, all eigenvalues of the Jacobian matrix for all defecting equilibrium are *x*_1_ and 0. Therefore, all defecting equilibrium is stable, as *−x*_1_ *<* 0. The mixed equilibrium is unstable, as the eigenvalue *x*_1_ is positive and it is always an eigenvalue in the mixed equilibrium as the first defector *j ≥* 2. All eigenvalues of the Jacobian matrix for the all cooperating equilibrium are *x*_*i*_ *− g*, (1 *≤ i ≤ n}*. Therefore, the all cooperating equilibrium is stable in the baseline system if max*{x*_*i*_, 1 *≤ i ≤ n} < g*, for all the 1 *≤ i ≤ n*. We consider the Prisoner’s dilemma situation in the baseline model, therefore, *x*_*i*_ *> g*. This indicate that all cooperation is unstable in the baseline.

For the defector sanction system, all eigenvalues of the Jacobian matrix for all defecting equilibrium is *ρf −x*_1_ and 0. Therefore, all defection equilibrium is stable in the defector sanction system if *ρf < x*_1_. The mixed equilibrium in the defector sanction system is stable if max(*x*_*i*_, *i < j*) *< ρf* and *x*_*j*_ *> ρf* when *j* is the first defector. All eigenvalues of the Jacobian matrix for the all cooperating equilibrium are *x*_*i*_ *− g − ρf* (1 *≤ i ≤ n*). Therefore, the all cooperating equilibrium is stable in the defector sanction system if *g* + *ρf >* max(*x*_*i*_, 1 *≤ i ≤ n*).

For the first role sanction system, All eigenvalues of Jacobian matrix for the all defecting equilibrium are *−x*_1_ and 0. Therefore, all defecting equilibrium is stable, as *−x*_1_ *<* 0. For the equilibrium to be a mixed equilibrium, *x*_1_ is always an eigenvalue when the first defector *j ≥* 2. As a result, the mixed equilibrium is unstable, as the eigenvalue *x*_1_ is positive. All eigenvalues of Jacobian matrix for the all cooperating equilibrium is *x*_*i*_ *− g*(2 *≤ i ≤ n*) and *x*_1_ *− g − f*. Therefore, the all cooperating equilibrium is stable in the 1st role sanction system if *g* + *f > x*_1_ and *g >* max(*x*_*i*_, 2 *≤ i ≤ n*).

